# Enhancing natural killer cell function with gp41-targeting bispecific antibodies to combat HIV infection

**DOI:** 10.1101/760280

**Authors:** Nitya S. Ramadoss, Nancy Q. Zhao, Barbra A. Richardson, Philip M. Grant, Peter S. Kim, Catherine A. Blish

**Affiliations:** Department of Biochemistry, School of Medicine, Stanford University, Stanford, CA, USA; Department of Medicine, Division of Infectious Diseases and Geographic Medicine, School of Medicine, Stanford University, Stanford, CA; Program in Immunology, School of Medicine, Stanford University, Stanford, CA; Department of Biostatistics, University of Washington, Seattle, WA, USA; Chan Zuckerberg Biohub, San Francisco, CA, USA; Stanford ChEM-H, Stanford University, Stanford, CA, USA

**Keywords:** HIV, NK cells, bispecific antibody, gp41, ADCC

## Abstract

**Objective(s):** To develop and evaluate the activity of bispecific antibodies (bsAbs) to enhance NK cell antibody-dependent cellular cytotoxicity (ADCC) against HIV-infected cells.

**Design:** These bsAbs are based on patient-derived antibodies targeting the conserved gp41 stump of HIV Env, and also incorporate a high affinity scFv targeting the activating receptor CD16 on NK cells. Overall, we expect the bsAbs to provide increased affinity and avidity over their corresponding monoclonal antibodies, allowing for improved ADCC activity against Env-expressing target cells.

**Methods:** bsAbs and their corresponding mAbs were expressed in 293T cells and purified. The binding of bsAbs and mAbs to their intended targets was determined using Bio-Layer Interferometry, as well as flow cytometry-based binding assays on *in vitro* infected cells. The ability of these bsAbs to improve NK cell activity against HIV-infected cells was tested using *in vitro* co-culture assays, using flow cytometry and calcein release to analyze NK cell degranulation and target cell killing, respectively.

**Results:** The bsAbs bound gp41 with similar affinity to their corresponding mAbs, and had increased affinity for CD16. The bsAbs also bound to primary CD4 T cells infected *in vitro* with two different strains of HIV. In addition, the bsAbs induce increased NK cell degranulation and killing of autologous HIV-infected CD4 T cells.

**Conclusions:** These bsAbs may provide a promising strategy to improve NK-mediated immune targeting of infected cells during HIV infection.

## Introduction

Natural killer (NK) cells are effector cells of the innate immune system that are equipped to recognize and rapidly eradicate tumor or virus-infected cells. Healthy cells escape NK cell killing because major histocompatibility complex class I (MHC-I) proteins on healthy cells engage NK cell inhibitory receptors [1]. In the context of HIV infection, downregulation of MHC-I proteins can activate NK cells; upregulation of stress-induced ligands on infected cells can trigger engagement of activating receptors on NK cells such as NKG2D [2]. HIV possesses mechanisms to counteract NK cell surveillance, such as downregulation of activating ligands [3, 4]. Another mechanism that impairs the ability of NK cells to control chronic HIV infection is the expansion of a dysfunctional CD56^−^CD16^+^ NK cell subset [5]. Thus, while NK cells have the potential to control HIV, enhancing their responses could optimize their ability to control HIV.

NK cell effector function can also be antibody-dependent - a process termed antibody dependent cellular cytotoxicity (ADCC). NK cells expressing the activating cell surface receptor FcγRIIIA (CD16) can bind antibodies via their constant (Fc) domain; these antibodies can in turn bind cell surface viral antigens. CD16-antibody binding activates NK cells, triggering the secretion of antiviral cytokines such as IFN-γ and the release of perforin and granzymes into target cells, resulting in cell death. ADCC may provide a protective or therapeutic benefit in HIV infection - ADCC antibody titers inversely correlate with viral load and rate of disease progression in HIV-infected individuals [6, 7], and are higher in elite controllers than in patients that progress to disease [8]. Furthermore, in mother-infant transmission of HIV, ADCC responses in breast milk correlate with reduced infection risk in infants via breastfeeding, while increasing survival rates in infected infants [9, 10]. In non-human primates, vaccine-induced ADCC antibody titers correlate with decreased viral loads and/or delayed disease progression after challenge with simian immunodeficiency virus (SIV) [11, 12]. In humans, the RV144 clinical trial, which was the first HIV vaccine trial to demonstrate some degree of efficacy [13], induced ADCC responses in vaccine recipients [14, 15]. Binding of IgGs to V1V2 of Env correlated inversely with the rate of infection, suggesting that ADCC-mediating antibodies could have contributed to the observed protection [16]. However, despite the evidence for the protective role of ADCC responses against HIV, it remains unclear whether the key driver of ADCC efficacy in infection is the binding to viral targets, the triggering of effector cell activation, or both.

Investigation into the epitopes of ADCC-mediating monoclonal antibodies (mAbs) in HIV have revealed multiple viral targets, including Env, Gag, Pol, Vpu, and Nef [17–21]. Within Env, many ADCC-mediating mAbs target gp120 [22]. Recently, however, a larger number of gp41-targeting mAbs that mediate ADCC have also been characterized [23, 24]. During viral membrane fusion, gp120 is shed, and gp41 undergoes a series of conformational changes that exposes various targetable epitopes such as the fusion peptide, the loop region that connects the two heptad regions, and the membrane proximal region [25]. The post-fusion state of gp41, a six-helix bundle [26–29], remains exposed at the cell surface after viral and host membranes fuse and is often referred to as a stump [30]. Stumps can also occur on cell surfaces as gp120 can be shed when virus buds from infected cells [30]. The cell-surface exposed nature of the stumps makes them targetable by anti-gp41, ADCC-mediating mAbs. However, unlike influenza or dengue virions that are densely studded with their envelope glycoproteins, HIV is sparsely coated with Env [31, 32]. This presumably results in very few targetable stumps per cell on the surfaces of infected cells, decreasing the overall efficacy of ADCC.

By engineering known anti-stump, ADCC-mediating mAbs into bispecific antibodies (bsAbs) that adopt a tetravalent IgG-scFv format [33], we rationalized that we would achieve the following. First, we would bridge NK cells to HIV-infected targets via CD16, a potent trigger of NK cell activation [34], thus forcing NK cells to contact and be activated specifically against these cells. Second, the bsAb would show increased avidity over a mAb due to its ability to simultaneously engage two CD16 receptors on the NK cell surface. The resulting increase in local concentration of bsAb at the NK cell surface can enable more efficient crosslinking to the stumps on the infected cell surface. As such, we tested these bsAbs for their ability to activate NK cells and clear HIV-infected primary CD4 T cells *in vitro.*

## Materials and methods

### Expression of bsAbs and mAbs

Monoclonal antibody (mAb) heavy and light chain plasmids were obtained from Dr. Julie Overbaugh (Fred Hutchinson Cancer Research Center). For the bispecific antibodies (bsAbs), the heavy chain sequence linked to the NM3E2 scFv sequence (IDT gBlocks) were cloned into the VRC01 heavy chain backbone by In-Fusion. NM3E2 scFv was expressed and purified from *E.coli* as described previously [35].

bsAbs and mAbs were expressed in Expi293 cells at a 2:1 Heavy:Light chain ratio, tracking viability (until <70%) or until day 6, whichever was sooner. Media was harvested and filtered, before purifying bsAbs and mAbs on Protein A columns, followed by size exclusion chromatography (S200 pg or S200 increase depending on scale, GE Healthcare). All proteins were analyzed by reducing and non-reducing SDS-PAGE.

### Antigens for BLI

6-helix containing a cysteine in the first HR2 helix (Ser68→Cys68) was expressed and purified as described previously [36]. The protein was reduced with immobilized TCEP resin (Pierce) using manufacturer’s protocols, and conjugated to EZ-Link™ Maleimide-PEG11-Biotin (Thermo Fisher). The extent of biotinylation was confirmed by HABA assay (Pierce). Excess biotin was removed by size exclusion. Commercial sources were used to obtain biotinylated CD16 (Acro Biosciences) and the biotinylated 19-mer adjacent loop peptides that overlap by 16 amino acids (Sigma). Loop peptide 1 is KDQQLLGIWGCSGKLICTT and loop peptide 2 is QLLGIWGCSGKLICTTAVP.

### Binding assays by BLI

Samples in assay buffer (1x TBS + 0.5% BSA) were dispensed into 96-well black flat-bottom plates (Greiner Bio-One) at a volume of 180–200 μl per well; all measurements were performed on the Octet Red96 (Pall ForteBio, Menlo Park, CA) at room temperature at 1000 rpm agitation. Streptavidin-coated biosensor tips (Pall ForteBio, Menlo Park, CA) were used to capture biotinylated antigens, and typical immobilization levels captured on the AMC sensors varied up to 0.5 nm. Antigen loading was carried out for 30s, followed by regeneration to remove non-specifically bound antigen. After two baseline steps of 20s each, association was performed for 60s in wells containing antibody dilutions, followed by dissociation in assay buffer for 90s. Between each antibody sample, a regeneration step was included to eliminate carryover. Raw data was exported and fitted using Prism’s (GraphPad Software) association then dissociation function.

### Binding assays by flow cytometry

CHO-WT was obtained through the NIH AIDS Reagent Program, from Dr. Carol Weiss and Dr. Judith White. These cells express envelope glycoprotein on their surface, are highly fusogenic and readily form syncytia with human CD4+ cells [37]. Cells were plated at 10^5^ cells/well in a 96-well flat-bottomed plate (Corning). After 24h, supernatant media was removed and replaced with FACS buffer. Cells were stained with 1:100 dilution of each antibody for 1h at 4°C, washed, and stained with Alexa 647-conjugated anti-human IgG (H+L) antibody (Thermo Fisher) for 1h at 4°C. Cells were further washed, trypsinized and analyzed by flow cytometry on the Accuri C6 (BD Biosciences).

For binding studies on HIV-infected primary human CD4 T cells, CD4 T cells were isolated from healthy human donor PBMCs by negative selection (Miltenyi Biotec), and activated with plate-bound anti-CD3 (clone OKT3, eBioscience), anti-CD28/CD49d (BD Biosciences) and PHA-L (eBioscience). After 48h, CD4 T cells were mock-infected, or infected with either Q23 [38] or NL4-3 (infectious molecular clone obtained through the NIH AIDS Reagent Program, from Dr. Malcom Martin [39]) virus, using Viromag magnetofection (OZ Biosciences). Both Q23 and NL4-3 viruses were grown in 293T cells and titrated as previously described [40]. 24h post infection, cells were stained at a 50nM concentration of each mAb/bsAb for 30 min at 4°C, washed, and then stained with Alexa 647-conjugated anti-human IgG (H+L) antibody (Thermo Fisher) for 30 min at 4°C. Cells were further washed, fixed with FACS Lyse (BD Biosciences) and analysed by flow cytometry on a MACSQuant Analyzer (Miltenyi Biotec).

### NK-CD4 co-culture assays

NK cells and CD4 T cells were isolated from donor PBMCs using magnetic activated cell sorting and negative selection (Miltenyi Biotec). NK cells were activated for 72h in 300IU/ml rhIL-2 (R&D Systems). CD4 T cells were simultaneously activated and infected as described above. NK cells were co-cultured with the infected or uninfected CD4 T cells at a 1:1 effector:target (E:T) ratio for 4h, either in the absence of or with increasing concentrations of bsAb or mAb. NK cells were identified by staining for CD3-V450 and CD56-PE-Cy7; their activation was monitored by staining for CD107a-APC (all antibodies from Biolegend). Cells were analyzed by flow cytometry on a MACSQuant Analyzer (Miltenyi Biotec). All flow cytometry analysis was performed using FlowJo version 9.9.6 (Tree Star).

For cytotoxicity assays, target CD4 T cells were loaded with 2.5uM calcein-AM (Invitrogen), and incubated with autologous IL-2 activated NK cells at a 4:1 E:T ratio. After a 4h incubation, cells were spun down and 100ul of culture supernatant was transferred into 96-well black flat-bottom plates (Greiner Bio-One), and calcein release into the supernatant was measured using a Synergy HTX fluorescent plate reader (Biotek), with control wells containing no effector cells (spontaneous lysis) and with 2% Tween-20 (maximum lysis). Specific killing was calculated using the equation: specific killing = (effector lysis - spontaneous lysis)/(maximum lysis - spontaneous lysis) x 100%.

### Statistical analysis

For co-culture assays, linear mixed effects models were used. The models incorporated fixed effects for concentration, bsAb vs mAb, and mock vs HIV infection, and a random effect for donor. This model takes into account that there are multiple observations per donor and accounts for variability between donors. Statistical analyses were performed using SPSS and the nlme package in R.

## Results

### Expression and purification of stump-targeting bsAbs

To improve ADCC activity of stump-targeting mAbs, we designed bsAbs in the IgG-scFv or Morrison format [33], in which the variable regions of the antibody target gp41 stumps on HIV-infected cells, while the high affinity scFvs tethered to the heavy chains bind CD16 on NK cells. These bsAbs can engage two CD16 receptors on the NK cell while binding to the target antigen, resulting in increased avidity. We chose the IgG-scFv format to retain full Fc effector function on the bsAb, so that we could directly compare bsAb activity to the corresponding mAb. We postulated that the combination of high affinity and avidity would drive high efficacy of the bsAbs compared to the mAbs, which have lower efficacy due to low-affinity Fcs engaging one CD16 on the NK cell surface at a time (Fig 1A).

**Figure 1.**
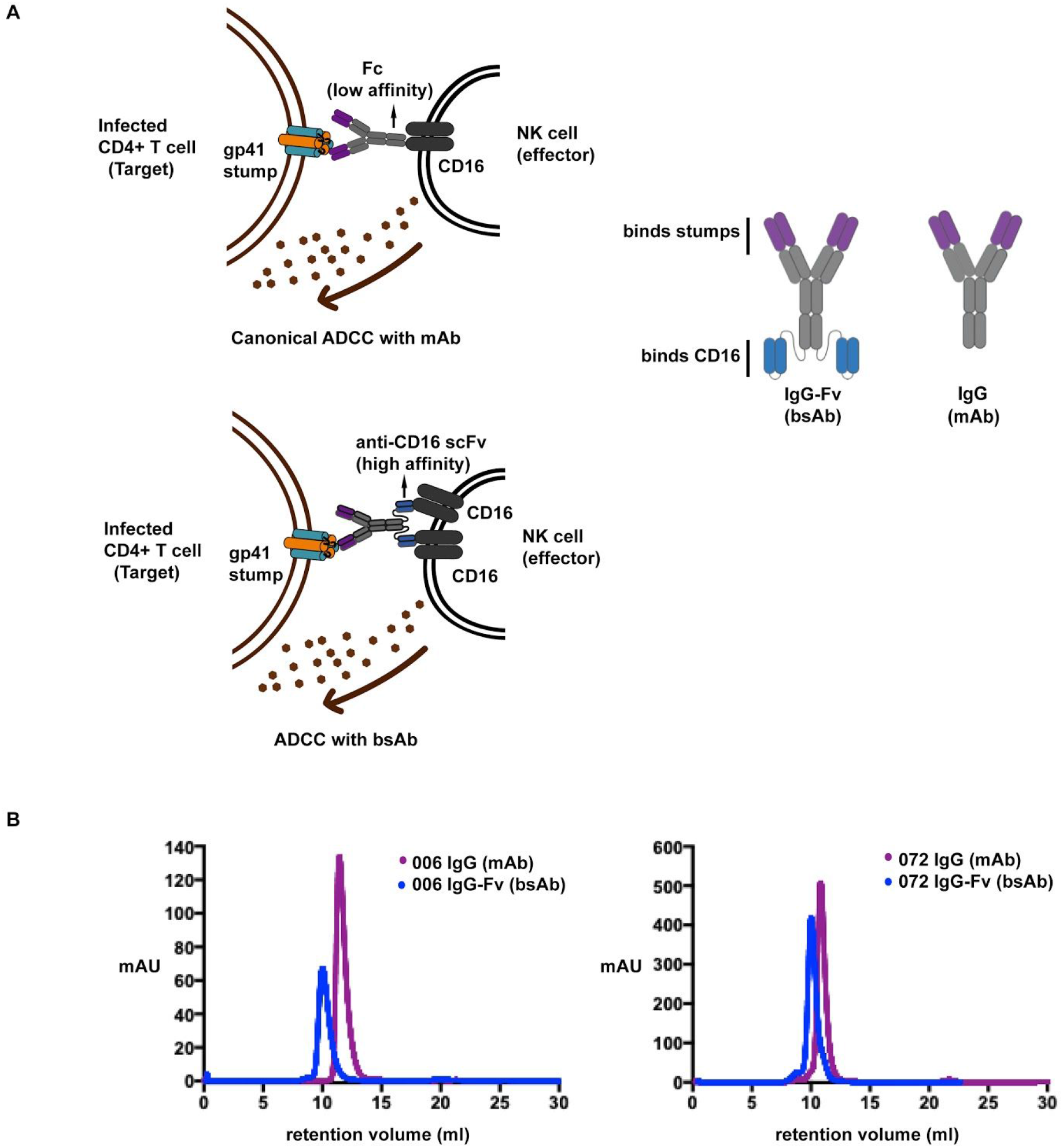
A. Schematic describing the difference between canonical ADCC with a mAb (top), and ADCC with a bsAb. A mAb binds to CD16 via its low affinity Fc to drive ADCC. With a bsAb, the high affinity of the scFvs tethered to the heavy chains, together with increased avidity from their binding two CD16 receptors on the NK cell potentially drives higher efficacy. This specific activation of NK cells results in lysis of the infected target via production of cytolytic granules. B. IgG-Fv bispecific antibodies (bsAbs) and corresponding stump-binding monoclonal antibodies (mAbs) were expressed in Expi293 cells, and purified by size exclusion chromatography on an S200 column after an initial protein A purification step. IgG-Fvs (~200kD, blue) elute earlier than IgGs (~150kD, purple) as expected.

For the stump binding antibodies, we used previously identified patient-derived anti-stump antibodies, QA255.006 and QA255.072, referred to hereafter as 006 and 072, respectively [24]. For the CD16-targeting scFv, we used NM3E2, a phage-display derived antibody [35]. After expression in mammalian cells, bsAbs and mAbs were isolated using Protein A resin, followed by size exclusion chromatography. As expected, the bsAbs eluted earlier than the mAbs on the size exclusion column due to their larger molecular weights (Fig 1B, Supplemental Fig 1).

### bsAbs show increased affinity to CD16 over mAbs, while engaging stumps with similar affinity

To determine whether bsAbs showed improved binding to CD16 over mAbs, we used bio-layer interferometry (BLI), and compared binding of the bsAbs, mAbs, and the NM3E2 scFv to CD16. The bsAbs demonstrated binding in the nanomolar range to CD16, which were 2-3 orders of magnitude improved over that of the mAbs (low to sub-micromolar range) (Fig 2A, Table 1). NM3E2 scFv affinity was determined to be 75.1 ± 15.1 nM, which is within an order of magnitude of the published value of 20 nM for this scFv [35].

**Figure 2.**
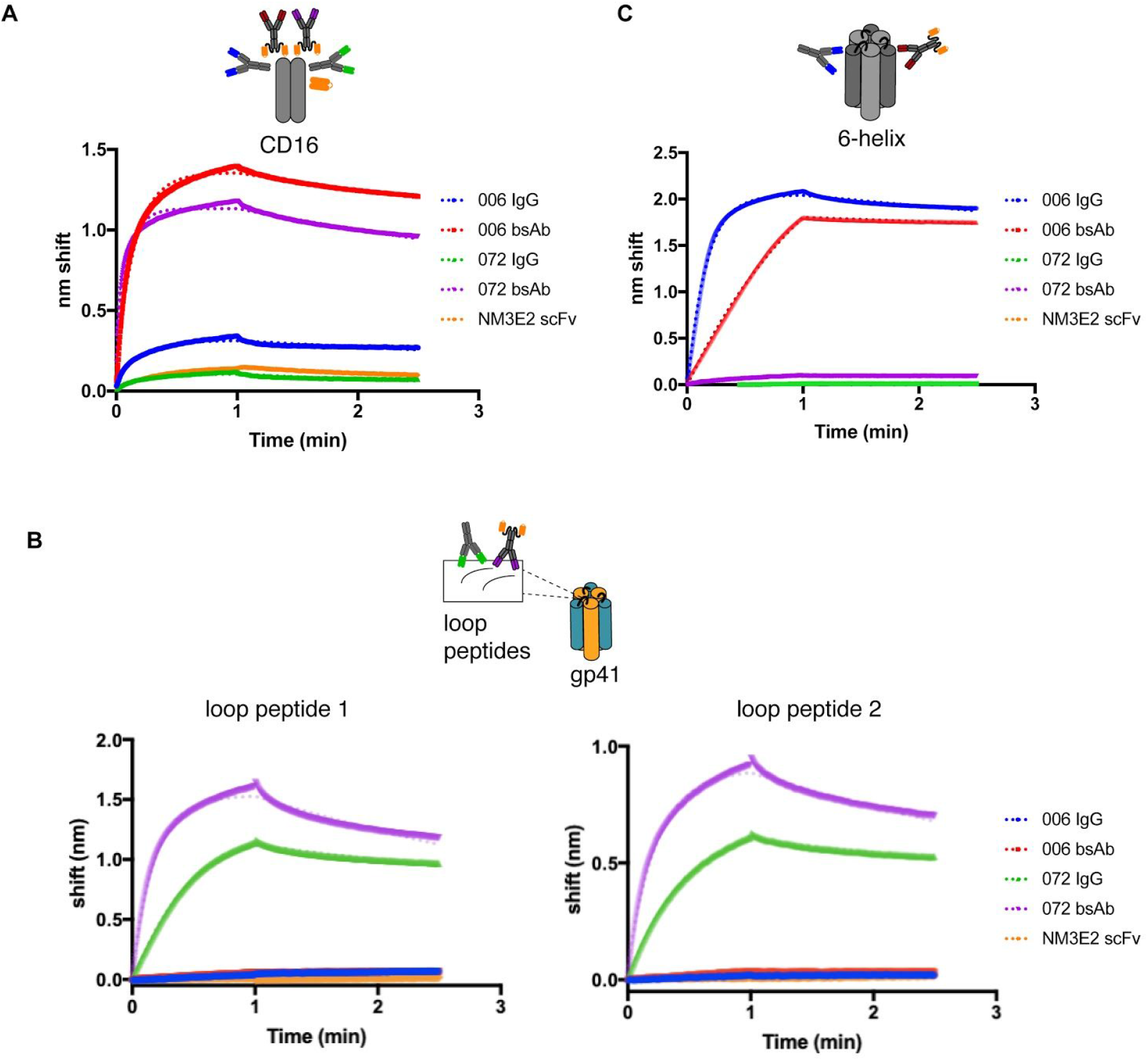
Binding of mAbs and bsAbs to, A) CD16 B) loop peptides 1 and 2 containing 072 epitope, and C) 6-helix as determined by Bio-layer Interferometry. Representative curves from 3 biological repeats shown. Fits are indicated in dotted line. Cartoon insets above each graph indicate the binding target tested in each case and the antibodies that are expected to bind.

**Table 1.** Apparent binding constants (K_app_) of Abs to 6-helix, loop peptides and CD16

|  | **006 IgG** | **006 IgG-Fv** | **072 IgG** | **072 IgG-Fv** | **NM3E2 scFv** |
| --- | --- | --- | --- | --- | --- |
| **CD16** | $0.37 \pm 0.26 \mu\text{M}$ | $4.6 \pm 2.4 \text{ nM}$ | $4.9 \pm 4.4 \mu\text{M}$ | $9.7 \pm 6.1 \text{ nM}$ | $72 \pm 14 \text{ nM}$ |
| **6-helix** | $2.5 \pm 0.55 \text{ nM}$ | $1.6 \pm 0.20 \text{ nM}$ | No binding | No binding | No binding |
| **Loop peptide 1** | No binding | No binding | $3.2 \pm 1.1 \text{ nM}$ | $2.5 \pm 0.67 \text{ nM}$ | No binding |
| **Loop peptide 2** | No binding | No binding | $3.1 \pm 1.4 \text{ nM}$ | $6.4 \pm 3.9 \text{ nM}$ | No binding |

To determine whether bsAbs retained their binding specificities and affinities to their targets, we used BLI and compared binding of the bsAbs, mAbs and NM3E2 to 6-helix, a stump mimetic that has been shown to be targeted by 006 but not 072 (since 6-helix does not contain the immunodominant loop that 072 binds to), as well as two 19-mer loop peptides that both contain the previously determined linear epitope of 072 [24], but that do not bind 006. Recombinantly expressed gp41 ectodomain was insoluble in the BLI assay buffer, and therefore not used in this assay. bsAbs and IgGs demonstrated similar levels of binding (Table 1), and retained their binding specificities to their corresponding targets, i.e., 006 bsAb and IgG bound 6-helix but not loop peptides, while 072 bsAb and IgG bound loop peptides but not 006 (Fig 2B, C).

### bsAbs bind to cells expressing Env, as well as to HIV-infected primary CD4+ T cells

The stump-binding mAbs have previously been shown to bind to Env in a cell-based ELISA assay [24]. To determine whether the constructs could engage their targets on the surface of cells, we first tested binding of the mAbs and bsAbs in adherent CHO cells expressing Env. mAbs and bsAbs bind 9-12% of Env-expressing CHO cells, 6-7 fold greater than the background secondary only control (Fig 3A). These results correlate with the binding of bsAbs and mAbs *in vitro,* showing that in the absence of NK cells or CD16, binding profiles of the bsAbs and mAbs to the target antigen are similar. We then tested dose-dependence of cell-surface binding with the 006 and 072 mAbs. Binding to Env-expressing CHO cells was dose-dependent for both 006 and 072 in the 0-500 nM range (Supplemental Figure 2A).

**Figure 3.**
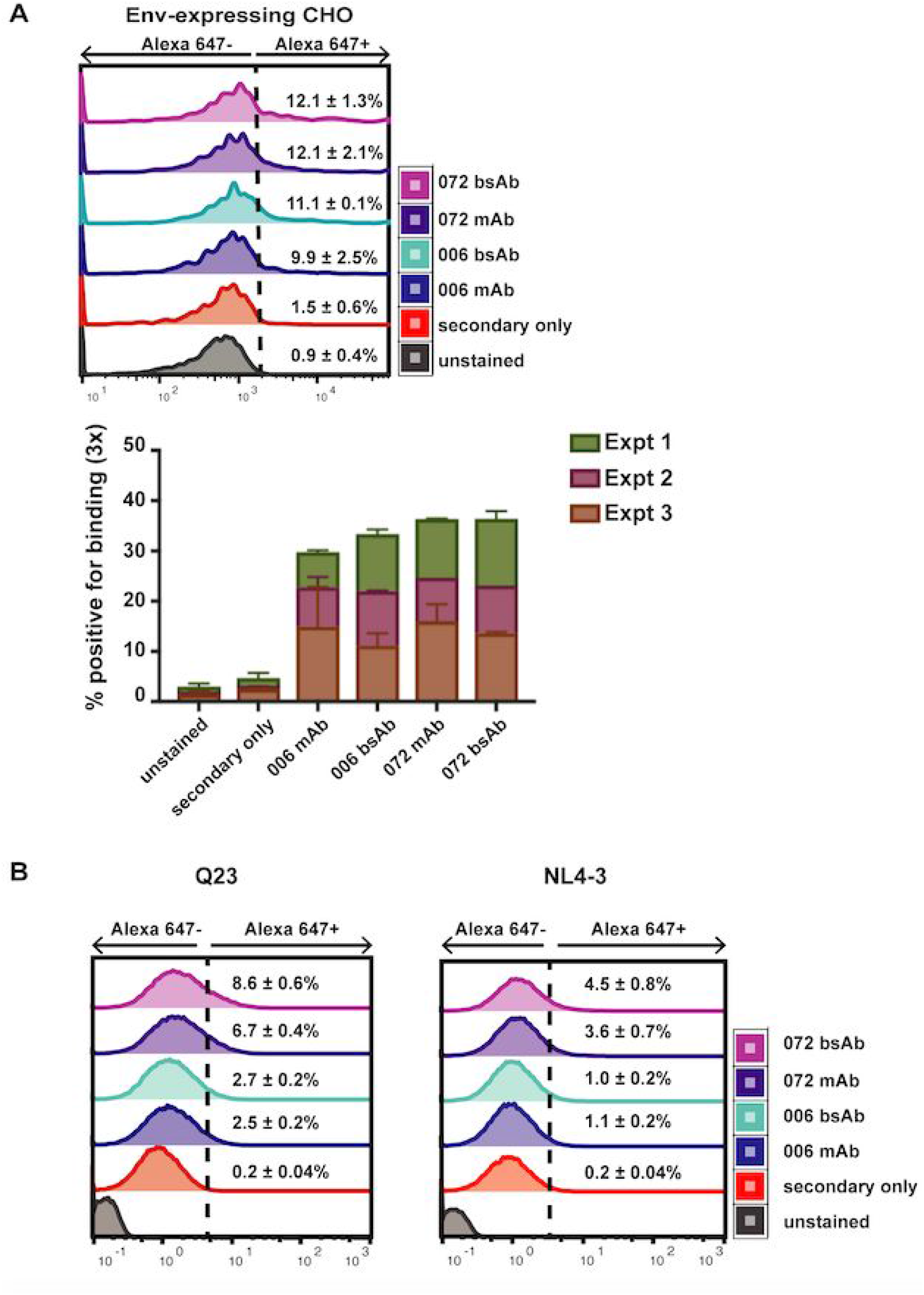
A. Binding of antibodies (bsAbs and mAbs) to CHO-WT cells expressing Env. Representative staggered histogram is shown for one of three biological repeats. Dotted line indicates gate set based on the secondary only-treated control. Percentage positive for antibody binding to the cell line for 3 biological repeats, and the corresponding SEM is indicated in stacked plot format. In the column graph, each segment represents an experiment set up in duplicate, and B. Binding of antibodies (bsAbs and mAbs) to primary CD4 T cells infected *in vitro* with either a clone from early subtype A HIV infection (Q23) or lab-adapted (NL4-3) strain of HIV. Histogram is shown for one representative repeat. Percentage positive for antibody binding for 6 biological repeats, and corresponding SEM is indicated.

To evaluate binding of these bsAbs to Env in a more physiological system, we tested the binding of the mAbs and bsAbs to primary CD4+ T cells infected *in vitro* with either a clone from early, subtype A infection (Q23) or a commonly used lab-adapted, subtype B strain (NL4-3). Both the 072 and 006 demonstrated binding to HIV-infected CD4 T cells, with 1-8% of infected cells staining positively; binding was similar between bsAbs and mAbs (Fig 3C). No binding higher than 1% was observed for either bsAb or mAb to mock-infected cells (Supplemental Figure 2B).

### bsAbs improve NK cell-mediated killing of autologous HIV-infected cells

To determine if the bsAbs could improve NK cell targeting of autologous HIV-infected cells, we used an *in vitro* NK-CD4 co-culture assay to test the effect of the bsAbs on NK cell degranulation and target cell killing. NK cell degranulation significantly increased, in co-culture with CD4 T cells infected with either the Q23 or NL4-3 strains, in the presence of increasing concentrations of the bsAb (Fig 4A). A similar increase was not observed with mock-infected cells. We also tested the 072 and 006 mAbs in the same assay, to determine if the bsAbs’ enhanced affinity and avidity improved NK cell targeting compared to the corresponding mAbs. As expected, the 072 and 006 bsAbs induced much stronger NK cell degranulation responses than the mAbs - at 10nM, NK degranulation was between 2.5 to 3.5-fold higher in the presence of the bsAbs compared to their matched mAbs (Fig 4A). For each of the 072 and 006 constructs, we used a linear mixed model [41], with a random effect for donor to account for inter-individual variability, to test the contributions of concentration, HIV infection and bsAb/mAb on NK cell degranulation. In the model, concentration, infection (with either strain of HIV) and bsAb (vs mAb) were all found to have a statistically significant effect on NK cell degranulation, for both bsAbs.

**Figure 4.**
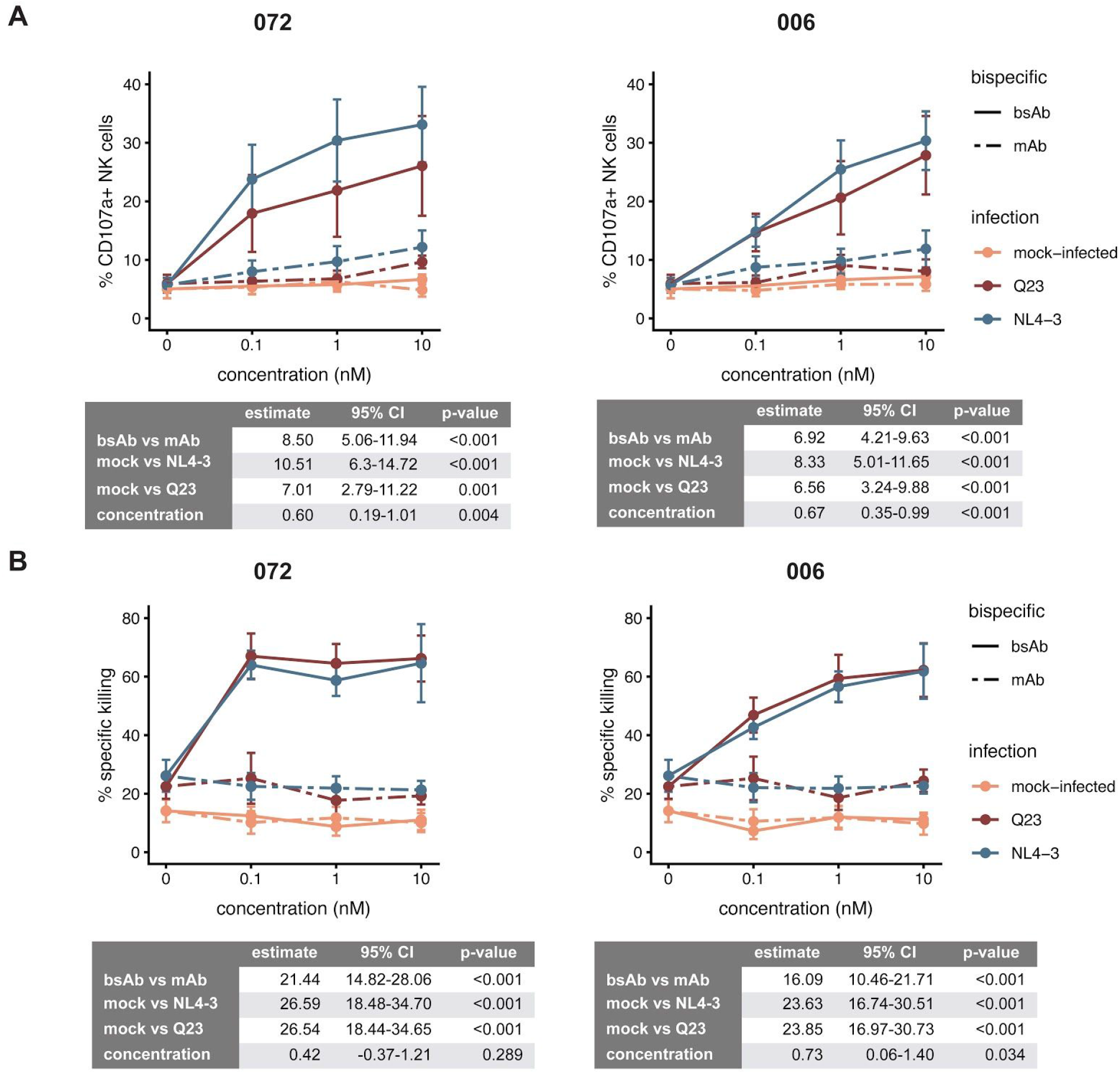
bsAbs increase NK cell responses to HIV-infected autologous CD4 T cells. Primary CD4 T cells were isolated from healthy donors, and mock-infected (orange) or infected with either an early subtype A clone (Q23, red) or a lab-adapted, subtype B (NL4-3, blue) strain of HIV. IL-2 activated NK cells from the same donor were co-cultured with these target cells, and NK cell responses in the presence of varying concentrations of the bsAbs/mAbs were assessed by A. degranulation of NK cells as measured by % positive for CD107a by FACS, and B. killing, as detected by calcein release. The data represents the mean and SEM for 6 (A) and 4 (B) donors respectively. Estimates for the contribution of each of the factors tested to % CD107a+ (A) or % specific killing (B) in the linear mixed model are shown in the table below each figure, together with 95% confidence intervals and p-value.

To determine if the increases in NK cell degranulation also led to increased killing of HIV-infected target cells, we measured target cell death using calcein release [42]. Both bsAbs significantly enhanced NK killing of HIV-infected cells with increasing concentration, for both strains of HIV tested, and had little nonspecific targeting, as killing of mock-infected cells was not increased (Fig 4B). Once again, the 072 and 006 bsAbs induced higher levels of target cell death in the presence of NK cells compared to their corresponding mAbs at matched concentrations (Fig 4B). In linear mixed models, infection (with either strain of HIV) and bsAb (vs mAb) were both found to have a statistically significant effect on NK cell degranulation; concentration had a statistically significant effect on the 006 bsAb but not the 072. The bsAb format enabled improved NK cell targeting and killing of infected cells over the mAb; this improved targeting was HIV-specific, and was observed across two HIV strains of different subtypes.

## Discussion

NK cells can rapidly recognize and lyse virus-infected or malignant cells; however, cancers and viruses have developed strategies to avoid NK cell surveillance. Thus, there is a need to specifically activate NK cells to overcome these evasion strategies. bsAbs have previously been used successfully in the cancer setting to enhance NK cell activity and clear tumors such as Non-Hodgkin’s lymphoma [43] and acute myeloid leukaemia [44]. In contrast, little work has been done in the field of bsAbs to recruit NK cells to treat viral infections. This work shows that NK cell activity against HIV can be specifically enhanced using bsAbs that simultaneously target gp41 on HIV-infected cells and the activating receptor CD16 on NK cells. As CD16 is expressed on other effector cell types such as macrophages and neutrophils [45, 46], activation of non-NK cells might be a concern; however, even if this were to occur, these effectors would likely augment the clearance of infected cells.

Recently, one study has linked single domain soluble CD4 to an anti-CD16 antibody to exploit the high affinity interaction between gp120 and its native receptor CD4 [47]. Soluble CD4 confers broad-spectrum recognition of envelope glycoprotein across different HIV strains. However, a potential limitation of this strategy is that CD4 can induce gp120 shedding, which can decrease binding over time. Our alternative approach of targeting gp41 avoids this limitation. Another abstract described NK cell activation in the context of HIV by linking a known gp120-binding broadly neutralizing antibody, VRC01 to an anti-CD16 single chain nanobody [48]. Our study differs from this aforementioned work as a proof of principle to determine the feasibility of using tetravalent bsAbs to enhance NK cell function against HIV, particularly by using gp41 antibodies to target infected cells. Presumably, this strategy can be extended to enhancing NK cell function in other disease contexts, such as influenza, where ADCC responses are also known to be protective [49].

The gp41 antibodies used in the bsAb constructs in this study were recently shown in a separate study to mediate ADCC in the commonly used RF-ADCC assay that employs cells coated with HIV Env [24]. In the context of *in vitro* infected cells, the binding of these mAbs was much lower (Fig 3C), which is unsurprising given the low density of Env on infected cell surfaces compared to protein-coated cell surfaces. Binding of the gp41 mAbs was increased when infecting cells with Vpu/Nef deficient virus (which results in higher Env surface expression), or in a cell based ELISA where soluble CD4 was used to induce gp120 shedding to expose the gp41 epitopes [24]. However, to overcome the reduced ADCC activity due to low binding of the mAbs in *in vitro* infected cells, we employed a more therapeutically translatable approach and enhanced ADCC activity by the tetravalent bsAb format to increase affinity and avidity to CD16, compared to the corresponding mAbs. The bsAb platform is an exciting tool to modulate NK cell function, with the potential for improved activity and specificity in future studies. For instance, studies targeting other activating receptors on NK cells, such as NKG2D, are warranted. Additionally, two distinct activating receptors on the NK cell can be targeted to engineer a trispecific antibody (two specificities to the NK cell, and one to the infected target) to fine tune NK activation and the magnitude of effector responses. Much of the focus for enhancing ADCC has previously only been on the level of the antibody - either on its Fab affinity to its target, or on its Fc type. Our work emphasizes the importance of improving effector cell activation to boost therapeutic outcomes.

Overall, we have demonstrated that NK cell cytotoxicity against HIV can be specifically enhanced using bsAbs targeting both the gp41 region of HIV Env and the activating receptor CD16 on NK cells, providing a new avenue for immune targeting of HIV infection.

## Acknowledgements

We are grateful for the gift of mAb plasmids from Dr. Julie Overbaugh at the Fred Hutchinson Cancer Research Center.

## Author contributions

NSR and NQZ designed and performed research, and analyzed data. BAR assisted in statistical analyses. PMG contributed samples to the study. PSK and CAB supervised the study. NSR, NQZ, PSK and CAB wrote the manuscript, with input from all authors.

---

**Figure S1.**
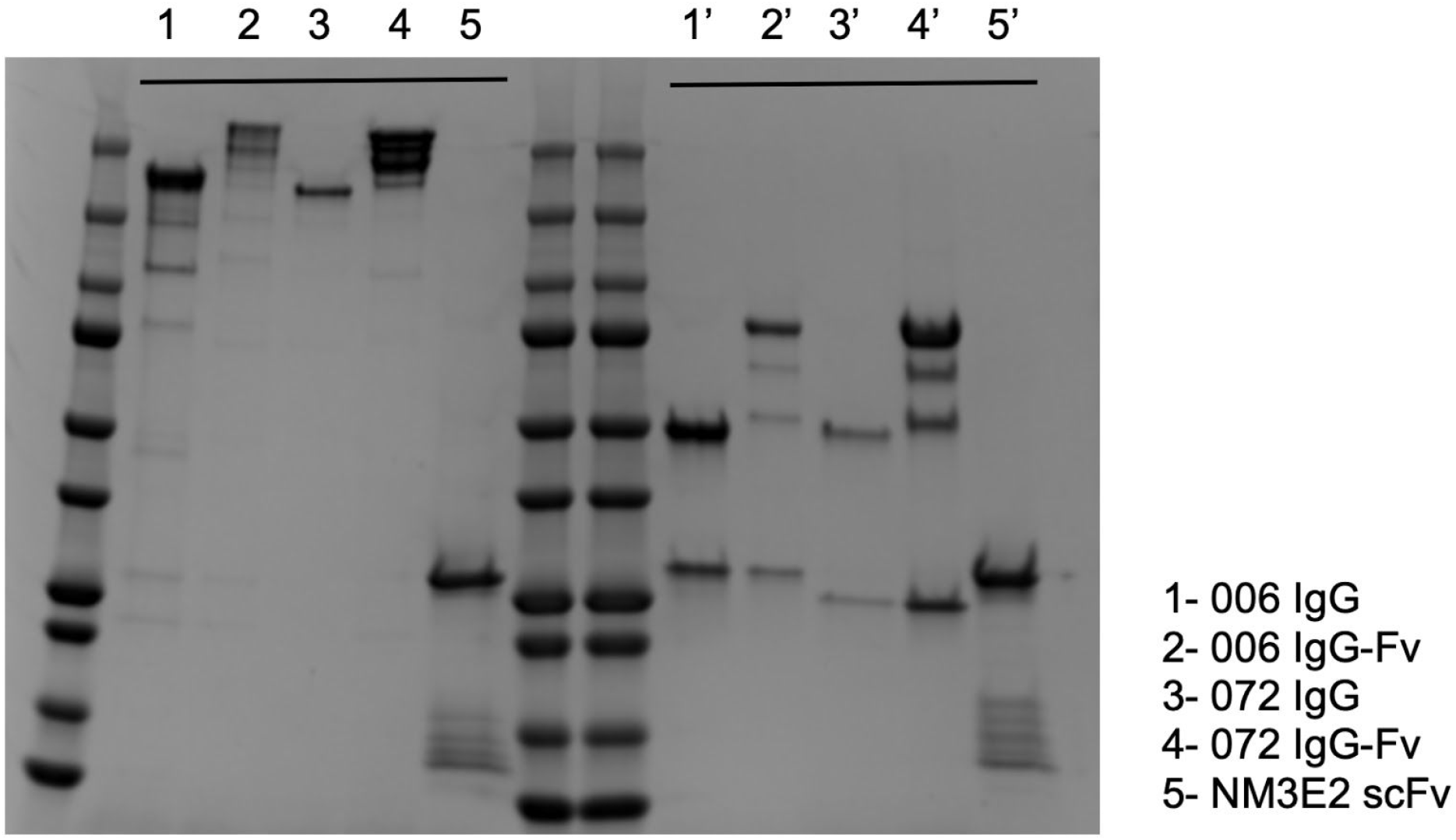
SDS-PAGE of bsAbs, mAbs and scFv post-expression under non-reducing and reducing (‘) conditions

**Figure S2.**
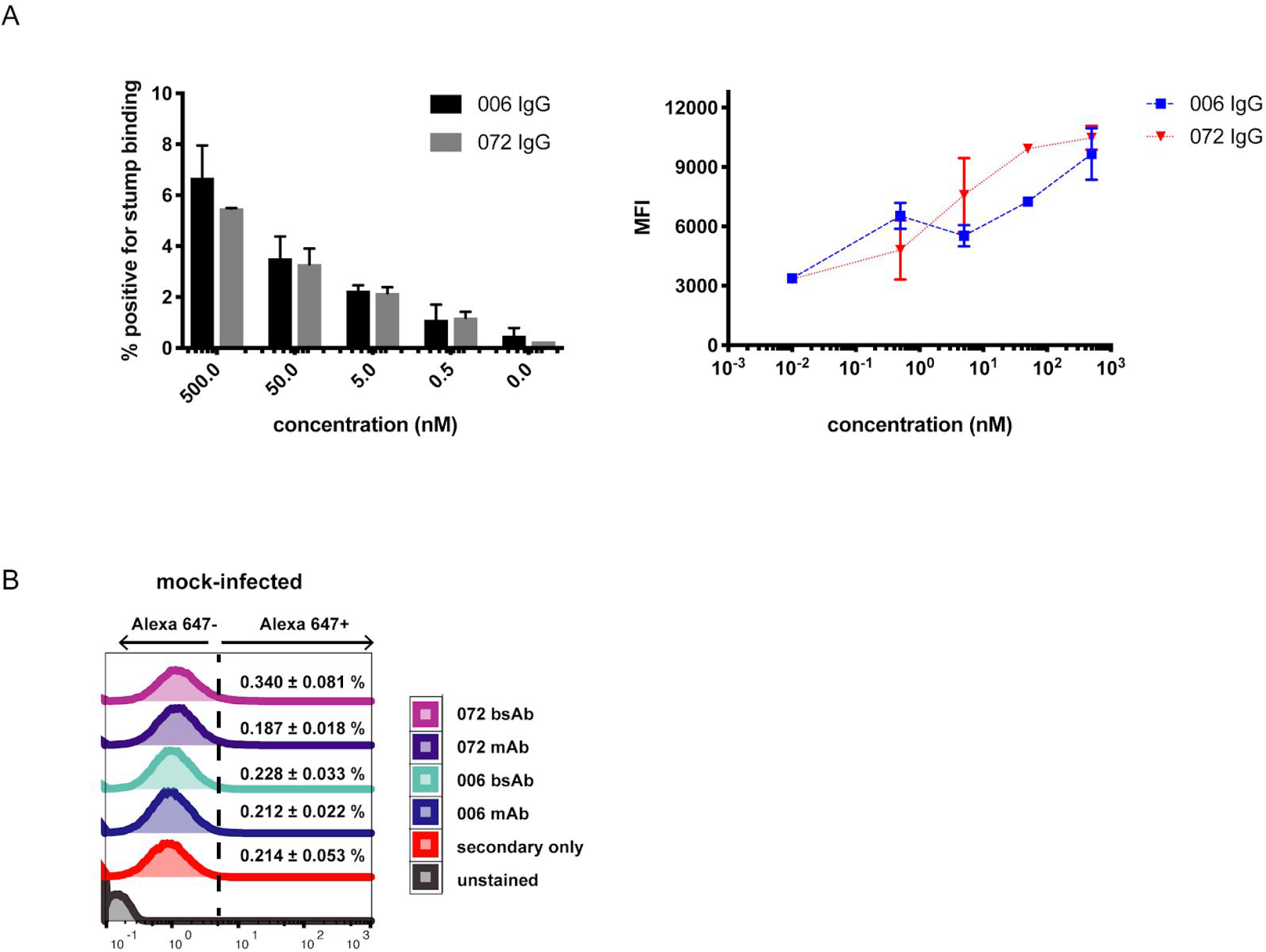
A. Dose-dependence of the binding of the 006 and 072 mAbs to CHO cells overexpressing Env, calculated as percent positive for stump binding (left) and by MFI (right). B. No binding of either 072 or 006 bsAbs or mAbs to mock-infected primary CD4 T cells.

## References

1 Pegram HJ, Andrews DM, Smyth MJ, Darcy PK, Kershaw MH. Activating and inhibitory receptors of natural killer cells. Immunol Cell Biol 2011; 89:216–224.

2 Richard J, Sindhu S, Pham TNQ, Belzile J-P, Cohen EA. HIV-1 Vpr up-regulates expression of ligands for the activating NKG2D receptor and promotes NK cell-mediated killing. Blood 2010; 115:1354–1363.

3 Cerboni C, Neri F, Casartelli N, Zingoni A, Cosman D, Rossi P, et al. Human immunodeficiency virus 1 Nef protein downmodulates the ligands of the activating receptor NKG2D and inhibits natural killer cell-mediated cytotoxicity. J Gen Virol 2007; 88:242–250.

4 Matusali G, Potesta M, Santoni A, Cerboni C, Doria M. The Human Immunodeficiency Virus Type 1 Nef and Vpu Proteins Down-regulate The Natural Killer Cell Activating Ligand PVR. J Virol 2012;: JVI–05788.

5 Mavilio D, Lombardo G, Benjamin J, Kim D, Follman D, Marcenaro E, et al. Characterization of CD56-/CD16+ natural killer (NK) cells: a highly dysfunctional NK subset expanded in HIV-infected viremic individuals. Proc Natl Acad Sci USA 2005; 102:2886–2891.

6 Ahmad R, Sindhu ST, Toma E, Morisset R, Vincelette J, Menezes J, et al. Evidence for a correlation between antibody-dependent cellular cytotoxicity-mediating anti-HIV-1 antibodies and prognostic predictors of HIV infection. J Clin Immunol 2001; 21:227–233.

7 Baum LL, Cassutt KJ, Knigge K, Khattri R, Margolick J, Rinaldo C, et al. HIV-1 gp120-specific antibody-dependent cell-mediated cytotoxicity correlates with rate of disease progression. J Immunol 1996; 157:2168–2173.

8 Lambotte O, Ferrari G, Moog C, Yates NL, Liao H-X, Parks RJ, et al. Heterogeneous neutralizing antibody and antibody-dependent cell cytotoxicity responses in HIV-1 elite controllers. AIDS 2009; 23:897–906.

9 Mabuka J, Nduati R, Odem-Davis K, Peterson D, Overbaugh J. HIV-specific antibodies capable of ADCC are common in breastmilk and are associated with reduced risk of transmission in women with high viral loads. PLoS Pathog 2012; 8:e1002739.

10 Milligan C, Richardson BA, John-Stewart G, Nduati R, Overbaugh J. Passively acquired antibody-dependent cellular cytotoxicity (ADCC) activity in HIV-infected infants is associated with reduced mortality. Cell Host Microbe 2015; 17:500–506.

11 Barouch DH, Liu J, Li H, Maxfield LF, Abbink P, Lynch DM, et al. Vaccine protection against acquisition of neutralization-resistant SIV challenges in rhesus monkeys. Nature 2012; 482:89–93.

12 Gómez-Román VR, Patterson LJ, Venzon D, Liewehr D, Aldrich K, Florese R, et al. Vaccine-elicited antibodies mediate antibody-dependent cellular cytotoxicity correlated with significantly reduced acute viremia in rhesus macaques challenged with SIVmac251. The Journal of Immunology 2005; 174:2185–2189.

13 Rerks-Ngarm S, Pitisuttithum P, Nitayaphan S, Kaewkungwal J, Chiu J, Paris R, et al. Vaccination with ALVAC and AIDSVAX to prevent HIV-1 infection in Thailand. N Engl J Med 2009; 361:2209–2220.

14 Karnasuta C, Paris RM, Cox JH, Nitayaphan S, Pitisuttithum P, Thongcharoen P, et al. Antibody-dependent cell-mediated cytotoxic responses in participants enrolled in a phase I/II ALVAC-HIV/AIDSVAX B/E prime-boost HIV-1 vaccine trial in Thailand. Vaccine 2005; 23:2522–2529.

15 Bonsignori M, Pollara J, Moody MA, Alpert MD, Chen X, Hwang K-K, et al. Antibody-dependent cellular cytotoxicity-mediating antibodies from an HIV-1 vaccine efficacy trial target multiple epitopes and preferentially use the VH1 gene family. J Virol 2012; 86:11521–11532.

16 Haynes BF, Gilbert PB, McElrath MJ, Zolla-Pazner S, Tomaras GD, Alam SM, et al. Immune-correlates analysis of an HIV-1 vaccine efficacy trial. N Engl J Med 2012; 366:1275–1286.

17 Grunow R, Franke L, Hinkula J, Wahren B, Fenyö EM, Jondal M, et al. Monoclonal antibodies to p24-core protein of HIV-1 mediate ADCC and inhibit virus spread in vitro. Clin Diagn Virol 1995; 3:221–231.

18 Isitman G, Stratov I, Kent SJ. Antibody-Dependent Cellular Cytotoxicity and NK Cell-Driven Immune Escape in HIV Infection: Implications for HIV Vaccine Development. Adv Virol 2012; 2012:637208.

19 Yamada T, Watanabe N, Nakamura T, Iwamoto A. Antibody-dependent cellular cytotoxicity via humoral immune epitope of Nef protein expressed on cell surface. J Immunol 2004; 172:2401–2406.

20 Wren LH, Chung AW, Isitman G, Kelleher AD, Parsons MS, Amin J, et al. Specific antibody-dependent cellular cytotoxicity responses associated with slow progression of HIV infection. Immunology 2013; 138:116–123.

21 Tiemessen CT, Shalekoff S, Meddows-Taylor S, Schramm DB, Papathanasopoulos MA, Gray GE, et al. Cutting Edge: Unusual NK cell responses to HIV-1 peptides are associated with protection against maternal-infant transmission of HIV-1. J Immunol 2009; 182:5914–5918.

22 Pollara J, Bonsignori M, Moody MA, Pazgier M, Haynes BF, Ferrari G. Epitope specificity of human immunodeficiency virus-1 antibody dependent cellular cytotoxicity [ADCC] responses. Curr HIV Res 2013; 11:378–387.

23 Yang Z, Liu X, Sun Z, Li J, Tan W, Yu W, et al. Identification of a HIV Gp41-Specific Human Monoclonal Antibody With Potent Antibody-Dependent Cellular Cytotoxicity. Front Immunol 2018; 9:2613.

24 Williams KL, Stumpf M, Naiman NE, Ding S, Garrett M, Gobillot T, et al. Identification of HIV gp41-specific antibodies that mediate killing of infected cells. PLoS Pathog 2019; 15:e1007572.

25 Harrison SC. Mechanism of Membrane Fusion by Viral Envelope Proteins. In: Advances in Virus Research.; 2005. pp. 231–261.

26 Chan DC, Fass D, Berger JM, Kim PS. Core structure of gp41 from the HIV envelope glycoprotein. Cell 1997; 89:263–273.

27 Weissenhorn W, Dessen A, Harrison SC, Skehel JJ, Wiley DC. Atomic structure of the ectodomain from HIV-1 gp41. Nature 1997; 387:426–430.

28 Lu M, Blacklow SC, Kim PS. A trimeric structural domain of the HIV-1 transmembrane glycoprotein. Nat Struct Biol 1995; 2:1075–1082.

29 Blacklow SC, Lu M, Kim PS. A trimeric subdomain of the simian immunodeficiency virus envelope glycoprotein. Biochemistry 1995; 34:14955–14962.

30 Moore PL, Crooks ET, Porter L, Zhu P, Cayanan CS, Grise H, et al. Nature of nonfunctional envelope proteins on the surface of human immunodeficiency virus type 1. J Virol 2006; 80:2515–2528.

31 Zhu P, Chertova E, Bess J Jr, Lifson JD, Arthur LO, Liu J, et al. Electron tomography analysis of envelope glycoprotein trimers on HIV and simian immunodeficiency virus virions. Proc Natl Acad Sci U SA 2003; 100:15812–15817.

32 Zhu P, Liu J, Bess J Jr, Chertova E, Lifson JD, Grisé H, et al. Distribution and three-dimensional structure of AIDS virus envelope spikes. Nature 2006; 441:847–852.

33 Coloma MJ, Morrison SL. Design and production of novel tetravalent bispecific antibodies. Nat Biotechnol 1997; 15:159–163.

34 Bryceson YT, March ME, Ljunggren H-G, Long EO. Synergy among receptors on resting NK cells for the activation of natural cytotoxicity and cytokine secretion. Blood 2006; 107:159–166.

35 McCall AM, Adams GP, Amoroso AR, Nielsen UB, Zhang L, Horak E, et al. Isolation and characterization of an anti-CD16 single-chain Fv fragment and construction of an anti-HER2/neu/anti-CD16 bispecific scFv that triggers CD16-dependent tumor cytolysis. Mol Immunol 1999; 36:433–445.

36 Root, M J, Kay, M S, Kim, P S. Protein Design of an HIV-1 Entry Inhibitor. Science. 2001; 291:884–888.

37 Weiss CD, White JM. Characterization of stable Chinese hamster ovary cells expressing wild-type, secreted, and glycosylphosphatidylinositol-anchored human immunodeficiency virus type 1 envelope glycoprotein. J Virol 1993; 67:7060–7066.

38 Poss M, Overbaugh J. Variants from the diverse virus population identified at seroconversion of a clade A human immunodeficiency virus type 1-infected woman have distinct biological properties. J Virol 1999; 73:5255–5264.

39 Adachi A, Gendelman HE, Koenig S, Folks T, Willey R, Rabson A, et al. Production of acquired immunodeficiency syndrome-associated retrovirus in human and nonhuman cells transfected with an infectious molecular clone. J Virol 1986; 59:284–291.

40 Strauss-Albee DM, Fukuyama J, Liang EC, Yao Y, Jarrell JA, Drake AL, et al. Human NK cell repertoire diversity reflects immune experience and correlates with viral susceptibility. Sci Transl Med 2015; 7:297ra115.

41 Bolker BM, Brooks ME, Clark CJ, Geange SW, Poulsen JR, Stevens MHH, et al. Generalized linear mixed models: a practical guide for ecology and evolution. Trends Ecol Evol 2009; 24:127–135.

42 Neri S, Mariani E, Meneghetti A, Cattini L, Facchini A. Calcein-acetyoxymethyl cytotoxicity assay: standardization of a method allowing additional analyses on recovered effector cells and supernatants. Clin Diagn Lab Immunol 2001; 8:1131–1135.

43 Glorius P, Baerenwaldt A, Kellner C, Staudinger M, Dechant M, Stauch M, et al. The novel tribody [(CD20)2xCD16] efficiently triggers effector cell-mediated lysis of malignant B cells. Leukemia 2012; 27:190.

44 Singer H, Kellner C, Lanig H, Aigner M, Stockmeyer B, Oduncu F, et al. Effective elimination of acute myeloid leukemic cells by recombinant bispecific antibody derivatives directed against CD33 and CD16. J Immunother 2010; 33:599–608.

45 Abeles RD, McPhail MJ, Sowter D, Antoniades CG, Vergis N, Manakkat Vijay GK, et al. CD14, CD16 and HLA-DR reliably identifies human monocytes and their subsets in the context of pathologically reduced HLA-DR expression by CD14hi/CD16neg monocytes: Expansion of CD14hi/CD16pos and contraction of CD14lo/CD16pos monocytes in acute liver fail. Cytometry A 2012; 81A:823–834.

46 Perussia B, Ravetch JV. FcγRIII (CD16) on human macrophages is a functional product of the FcγRIII-2 gene. Eur J Immunol 1991; 21:425–429.

47 Li W, Wu Y, Kong D, Yang H, Wang Y, Shao J, et al. One-domain CD4 Fused to Human Anti-CD16 Antibody Domain Mediates Effective Killing of HIV-1-Infected Cells. Sci Rep 2017; 7:9130.

48 Davis ZB, Lenvik T, Hansen L, Felices M, Cooley S, Vallera D, et al. A Novel HIV Envelope Bi-Specific Killer Engager Enhances Natural Killer Cell Mediated ADCC Responses Against HIV-Infected Cells. Blood 2016; 128:2517–2517.

49 Jegaskanda S, Weinfurter JT, Friedrich TC, Kent SJ. Antibody-dependent cellular cytotoxicity is associated with control of pandemic H1N1 influenza virus infection of macaques. J Virol 2013; 87:5512–5522.

